# Accounting for body mass effects in the estimation of field metabolic rates from body acceleration

**DOI:** 10.1101/2020.07.24.219204

**Authors:** Evan E. Byrnes, Karissa O. Lear, Lauran R. Brewster, Nicholas M. Whitney, Matthew J. Smukall, Nicola J. Armstrong, Adrian C. Gleiss

**Affiliations:** Centre for Sustainable Aquatic Ecosystems, Harry Butler Institute, Murdoch University, 90 South St. Murdoch, WA 6150 AUS; College of Science, Health, Engineering and Education, Murdoch University, 90 South St. Murdoch, WA 6150 AUS; Bimini Biological Field Station Foundation, South Bimini, BAH; Harbor Branch Oceanographic Institute, Florida Atlantic University, 5600 N US Highway 1, Fort Pierce, FL, 34946 USA; Anderson Chabot Centre for Ocean Life, New England Aquarium, 1 Central Wharf, Boston, MA, 02110 USA; College of Fisheries and Ocean Sciences, University of Alaska Fairbanks, 2150 Koyukuk Dr. Fairbanks, AK 99775 USA; Mathematics and Statistics, Murdoch University, 90 South St. Murdoch, WA, Australia

**Author notes:** Address correspondence to E.E. Byrnes.

**Keywords:** acceleration, biologging, ecophysiology, elasmobranch, field metabolic rate, respirometry

## Abstract

Life history, reproduction, and survival are fundamentally linked to energy expenditure and acquisition. Dynamic Body Acceleration (DBA), measured through animal-attached data-loggers or transmitters, has emerged as a powerful method for estimating field metabolic rates of free-ranging individuals. After using respirometry to calibrate oxygen consumption rate 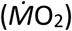 with DBA in captive settings, predictive models can be applied to DBA data collected from free-ranging individuals. However, laboratory calibrations are generally performed on a narrow size range of animals, which may introduce biases when predictive models are applied to differently sized individuals in the field. Here, we tested the influence of scale effects on the ability of a single predictive model to predict 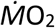 over a range of body sizes. We performed respirometry experiments with individuals spanning one order of magnitude in body mass (1.74–17.15 kg) and used a two-step modelling process to assess the intra-specific scale dependence of the 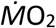-DBA relationship and incorporate such dependencies into the covariates of 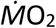 predictive models. The final predictive model showed scale dependence; the slope of the 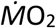-DBA relationship was strongly allometric (M^1.55^), whereas the intercept term scaled closer to isometry (M^1.08^). Using bootstrapping and simulations, we tested the performance of this covariate-corrected model against commonly used methods of accounting for mass effects on the 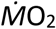-DBA relationship and found lowest error and bias in the covariate-corrected approach. The strong scale dependence of the 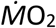-DBA relationship indicates that caution must be exercised when models developed using one size class are applied to individuals of different sizes.

**Summary statement:** The relationship between oxygen consumption rate and dynamic body acceleration is allometrically dependent, and models incorporate different slope and intercept scaling rates estimate metabolic rates more accurately than mass-specific approaches.

## INTRODUCTION

Energy is perhaps the most essential resource required by animals, and fundamentally governs their life history, behaviour, and ecology (Brown et al., 2004; Marquet et al., 2004). Once basal energy requirements are met, remaining energy can be allocated to various behaviours, growth, or reproduction. As such, rates at which animals acquire and expend energy and how excess energy is partitioned between different tasks directly impacts their survival and fitness (Jobling, 1993; Nisbet et al., 2012). Therefore, being able to estimate the energy demands of free-ranging animals, i.e. field metabolic rates (FMR) is fundamental to understanding their biology, behaviour, and influence on wider ecosystem and population processes.

Measuring the metabolic rates of animals in the laboratory is a well-established practice using either direct or indirect calorimetry, however, estimating FMR is more challenging. Commonly used methods to estimate FMR include measuring CO_2_ production via doubly labelled water (DLW; Speakman, 1997) or measuring heart rate (*f*_*H*_) or dynamic body acceleration (DBA) as a proxy for oxygen consumption rate 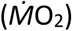. Of these, albeit having its own limitations (reviewed in Wilson *et al.* 2020; but see Gleiss, Wilson, Shepard 2011), the DBA technique has come to the fore because of its wide taxonomic applicability, high temporal resolution (in contrast to DLW; Butler et al., 2004), and the logistical simplicity of attaching acceleration data-loggers, as opposed to the invasive surgery required for implantation of heart-rate loggers (Green et al., 2009). However, before using DBA to predict FMR, laboratory calibrations are necessary to establish predictive models that relate DBA to 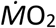. Such calibrations have been conducted for numerous vertebrate and invertebrate species and show that a linear relationship between 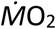 and DBA holds across all taxa, where the intercept constitutes the standard or basal metabolic rate (SMR/BMR) and the slope indicates how 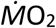 changes with activity (Wilson et al., 2020). After calibrations, predictive models can then be applied to field-measured DBA data to predict the FMR of free-ranging animals.

To discern the utility of any given FMR estimation method, it is necessary to test its sensitivity to factors that are expected to influence 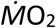 and to incorporate appropriate corrections into estimations. For instance, temperature and body size are considered the most important factors that influence metabolism, among other factors (e.g. age, sexual maturity, specific dynamic action), with both displaying positive exponential effects on 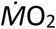 in ectotherms (Clarke and Johnston, 1999; Gillooly et al., 2001). As temperature is easily manipulated under laboratory conditions, the temperature dependence of the 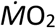-DBA relationship has been tested by numerous studies. As such, it is known that the intercept of the 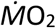-DBA relationship changes with temperature according to a van’t Hoff or Arrhenius relationship, while the slope is unaffected (e.g. Lear et al., 2017; Lyons et al., 2013). Accordingly, temperature variation can be accounted for by incorporating a temperature-dependent intercept term into predictive 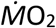 models. Although the level of temperature dependence varies between species, this type of correction has proven to be effective across ectothermic taxa, including fish (2020; Lear et al., 2017; Wright et al., 2014), reptiles (Enstipp et al., 2011), crustaceans (Lyons et al., 2013), and molluscs (Robson et al., 2016).

Unlike temperature, it is unknown how body size affects the 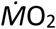-DBA relationship, due to difficulties with holding large species in captivity (Whitney et al., 2018). However, it is imperative that body mass effects are incorporated into estimations of 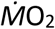, given the well-established allometric scaling patterns of SMR/BMR (da Silva et al., 2006; Kleiber, 1932). Defined by a power law: 

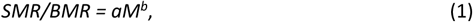

the SMR/BMR of organisms increases positively with body mass (*M*) at a rate defined by a species specific scaling coefficient (*a*) and exponent (*b*) (White et al., 2007). Thus, given that the intercept of the 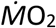-DBA relationship is representative of SMR/BMR, it is expected that the 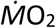-DBA intercept should scale allometrically with body mass.

While there is a reasonably well-defined hypothesis for the allometry of SMR/BMR, the kinematics that define animal movement and acceleration, and therefore govern the slope of the 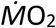-DBA relationship, adhere to a different set of allometric laws. For example, physiological forces that govern variations in SMR/BMR relate to factors such as fractal oxygen-transport networks and body surface area to volume ratios, which control the supply of metabolic substrates, and scale at allometric rates near 0.75 (West et al., 1997) and 0.67, respectively (Kozłowski et al., 2003; West and Brown, 2005). In contrast, DBA is a product of body kinematics, such as stride frequency and amplitude, which scale at allometric rates of approximately -0.33 (Bale et al., 2014) and 0.33 (Videler, 1993), respectively. Thus, it would be prudent to assume that the allometry of the 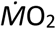-DBA slope, which represents changes in oxygen consumption rates in response to changes in physical activity, is not the same as the intercept. Nonetheless, current methods account for the effect of body mass by applying either isometric (Bouyoucos et al., 2017; O2 kg-1, e.g. Payne et al., 2011) or allometric (O_2_ kg^-b^, e.g. Enstipp *et al.* 2011; Lear *et al.* 2020) mass corrections to 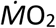 estimates, where *b* is a species specific SMR/BMR scaling exponent. These corrections assume an identical slope and intercept mass-scaling. However, if the 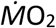-DBA slope was to scale at a different rate than SMR/BMR, it is likely that these methods introduce significant bias into 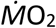 estimates.

The present study tested for allometric scaling rates in both the intercept and slope of the 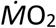-DBA relationship and develops an approach for incorporating mass effects into 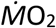 predictive models. To do this, we conducted respirometry experiments on individuals spanning one order of magnitude in body mass (1.74–17.15 kg), using lemon sharks (*Negaprion brevirostris*) as a model species. We then used a two-step modelling process to test for independent mass-scaling of the intercept and slope of the 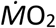-DBA relationship and incorporated the established mass-scaling effects into 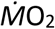 predictive models. To compare our approach against other established models, we ran a series of simulations to assess model bias and error.

## MATERIALS AND METHODS

### Capture and maintenance

Respirometry experiments were conducted on two groups of lemon sharks. The first group consisted of individuals <3 kg in mass (n=16, 69.5–86.0 cm total body length [TL]), captured off Florida, USA and housed at Mote Marine Laboratory (MML) in Sarasota, FL, USA for the duration of experiments. The second group consisted of individuals >3 kg in mass (n=5, 107.0–154.0 cm TL), captured and housed off South Bimini, BHS near Bimini Biological Field Station (BBFS). All sharks were fasted for at least 48 hours prior to experiments to ensure a post-absorptive state and allow them to recover from capture stress. Further capture and maintenance details for are provided in Lear et al. (2017) (MML) and the supplementary information (BBFS).

### Respirometry trial protocol

For MML sharks, respirometry was conducted between 2015–2016 as part of a separate study (Lear et al., 2017). A closed, annular, static respirometry system was constructed from a modified 2.45 m diameter fibreglass holding tank, as described in Whitney et al. (2016). Sharks were acclimated to the system for 12 hours prior to trials. Trials began near 100% air saturation and were run until dissolved oxygen (DO) levels reached 80% air saturation. DO (% air saturation and mg L^-1^) and water temperature (°C) were measured by a handheld multiparameter meter (Pro Plus, Yellow Springs Instruments, Yellow Springs, OH, USA) and recorded by researchers every five minutes throughout trials. To assess background respiration, a blank respirometer was measured for four hours following each set of trials. For full details of MML trial protocol, see Lear et al. (2017).

BBFS trials were conducted using a field-based respirometry system, similar to Byrnes et al. (2020). Two closed, annular, static respirometry systems were constructed from modified polyvinyl-lined metal-frame pools (Bestway Corp., London, UK), which were erected on a levelled section of beach on South Bimini, BHS. A 2.44 m diameter pool was used for sharks <110 cm TL, whereas a 3.66 m diameter pool was used for sharks >110 cm TL. Sharks were transferred into the system eight hours prior to trials to allow them to acclimate. Trials began with air saturation between 80 and 110% and were run until saturation decreased by 20 percentage points or reached 70% saturation, whichever occurred first. Throughout trials, DO (% air saturation and mg L^-1^) and water temperature (°C) were measured and recorded by a multiparameter meter (HI98196, Hanna Instruments, RI, USA) every 30 seconds. To assess background respiration, a blank respirometer was measured for at least 90 minutes immediately following the final trial for each shark. Further details on BBFS trials are provided in supplementary information.

Throughout all MML and BBFS trials, sharks were equipped with a Cefas G6a+ acceleration data logger (Cefas Inc., Lowestoft, UK), mounted through the base of the first dorsal fin as per Lear *et al.* (2017), which continuously recorded triaxial acceleration at 25 Hz.

### Body acceleration and oxygen consumption rate calculation

Vectorial dynamic body acceleration (VeDBA), a measurement of physical activity, was calculated from acceleration data recorded during respirometry. VeDBA was chosen as the acceleration metric for modelling instead of the commonly used overall dynamic body acceleration (ODBA) to allow predictive models to be applied to data collected by acceleration transmitters, which process acceleration onboard as VeDBA. Additionally, VeDBA produces more robust estimations of activity when tag orientation may differ between applications, such as external mounting of loggers versus internal implantation of transmitters (Qasem et al., 2012).

To match the sampling frequency of acceleration transmitters, which record long-term field data at 5 Hz, raw acceleration data were down-sampled by decimation from 25 Hz to 5 Hz, which has been shown to be a sufficient sampling frequency for estimating oxygen consumption rates of fishes (Brownscombe et al., 2018). Gravitational acceleration was estimated by calculating a 4s running mean, which was then subtracted from the raw acceleration, for each channel, to derive dynamic body acceleration (DBA). VeDBA was calculated as the vectorial sum of these DBA derivations: 

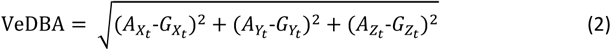

where 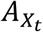, 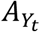, and 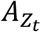 are the raw acceleration values and 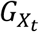, 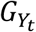, 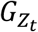 are the gravitational acceleration observed for each channel at time *t*. Acceleration data were processed using Igor Pro (Version 7.08; Wavemetrics, Lake Oswego, OR, USA) and the Ethographer extension (Sakamoto et al., 2009).

Dynamic acceleration data were visually examined to identify intervals where sharks maintained consistent behaviour (either resting or swimming) for 15–20 minutes during trials. Higher sampling frequencies of DO used during BBFS than MML trials allowed for shorter calibration intervals to be used for BBFS trials. Therefore, intervals of at least 20 minutes were used for MML sharks (Lear et al., 2017), whereas intervals of 15 minutes were used for BBFS sharks to increase sample sizes for BBFS trials. Mean water temperature, VeDBA and whole animal oxygen consumption rate (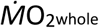; mg O_2_ hr^-1^) were calculated for each interval. 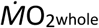 was calculated using the following equation from measurements of DO within respirometers: 

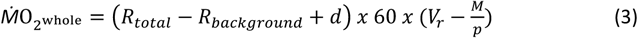

where *R*_*total*_ is rate of decline of DO during the calibration interval in mg O_2_ L^-1^ min^-1^ (estimated using linear regression), *R*_*background*_ is the rate of background respiration in mg O_2_ L^-1^ min^-1^, *d* is the diffusion rate of oxygen into the system (0.0002 mg 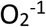 L^-1^ min^-1^; Byrnes et al. 2020), V_r_ is the respirometer volume in L, *M* is the mass of the shark in kg, and *p* is the density of the shark in kg L^-1^. All sharks were assumed to have a density of 1.0556 kg L^-1^, the average density reported for adult *N. brevirostris* in Florida (Baldridge Jr, 1970).

To remove variation in oxygen consumption rates due to temperature, all 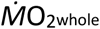 estimations were temperature corrected to 29.5°C, the mean water temperature of MML trials using: 

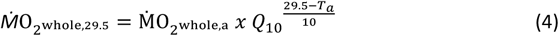

where 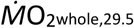 is the 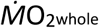 calculated at temperature 29.5°C, 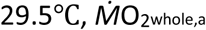 is the 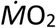 whole calculated at temperature *T*_*a*_, and *Q*_*10*_ is the temperature scaling factor for lemon sharks. A *Q*_*10*_ of 2.96 was used to correct 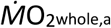 during resting intervals, and a *Q*_*10*_ of 1.69 was used to correct 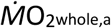 during active intervals (Lear et al., 2017).

Prior to building predictive models, VeDBA estimates were corrected for measurement error due to sensor noise by subtracting the mean VeDBA from resting intervals. Note, while baselining VeDBA as such may result in some minorly negative values, this correction was necessary to ensure that the model intercept represented SMR of sharks because measurements from acceleration data loggers record small acceleration values even while at complete rest (Gleiss et al., 2010). Statistical significance was declared at *p*≤0.05 and all statistical analysis was carried out in R v3.6.3 (R Core Team, 2019).

### Model development: Coefficient-corrected Approach

As previously discussed, the relationship between metabolic rate and activity (the 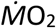-VeDBA relationship) is known to be linear (Wilson *et al.* 2020), described by the equation: 

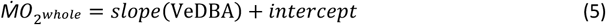

However, the slope and intercept theoretically follow allometric relationships, according to the power law defined in Equation 1. As such, a model which incorporates allometric scaling corrections into the slope and intercept terms would adhere to the following equation: 

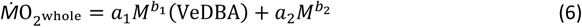

where *M* is the mass of the animal, *a*_*1*_ and *a*_*2*_ are the scaling coefficients for the slope and intercept terms, respectively, and *b*_*1*_ and *b*_*2*_ are the scaling exponents for the slope and intercept terms, respectively. To estimate such a model, we developed a two-step modelling process, similar to Halsey *et al.* (2009). In the first step (section 2.3.2), we used a linear mixed-effects model (LME) to test the mass dependency of the slope and intercept in Equation 5 (Fig. 1). In the second step (section 2.3.3), the slope and intercept terms from the best-fit LME were extracted and used to determine the allometry of each term (Fig. 1). The accuracy of the final predictive model formed by this process was error tested and compared against previous methods for correcting 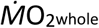 estimations for body mass (section 2.4).

**Fig. 1.**
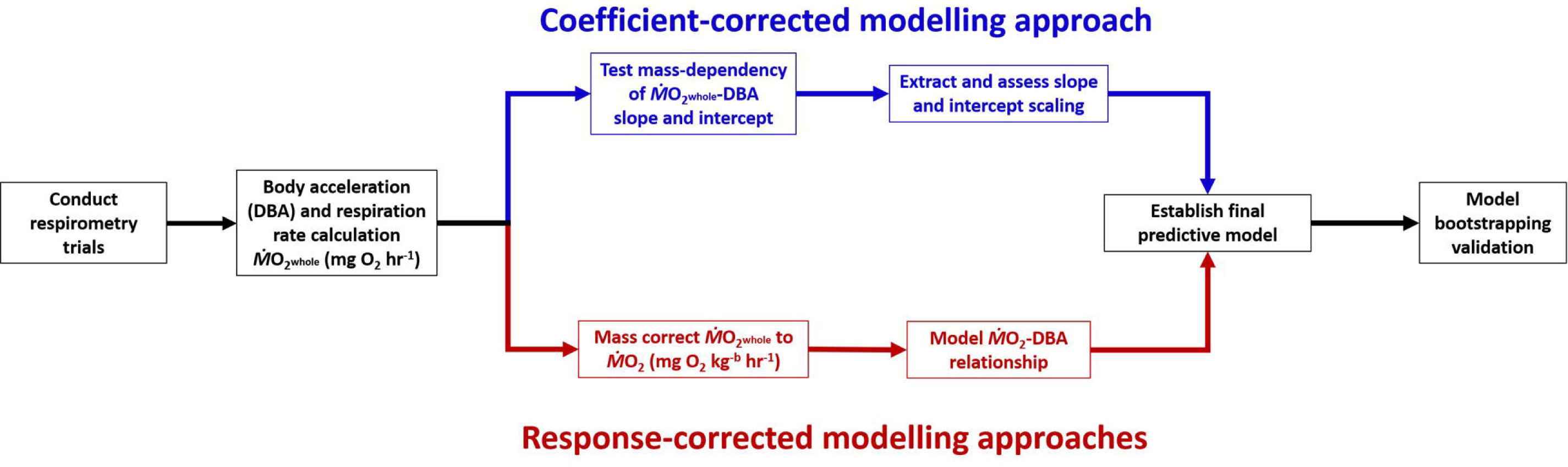
Workflow of establishing 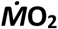 predictive models from respirometry and body acceleration data. All steps from initial data collection through final model validation are shown for the covariate-corrected modelling approach (established herein; top) and commonly used response-corrected approaches (bottom). Mass-corrections in response-corrected approaches can be either isometric (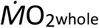 divided by M^1^) or allometric (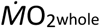 divided by M^b^), where b is a species-specific scaling rate.

### Testing mass-dependency of slope and intercept

Due to sharks frequently switching between inactive and active behaviours in MML trials, 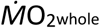 observations were not available over a sufficient range of acceleration values to produce a linear fit for each individual (Table S1). To increase the number of 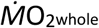 observations and range of sampled activity levels for each body mass, MML sharks were grouped into 400 g mass classes (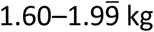, 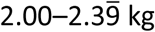, 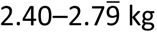, and 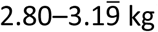). The midrange of a class was used as the mass for all sharks within the class during modelling. The BBFS sharks, on the other hand, showed a greater range of activity within individual trials and had a larger range in body size, and were therefore kept as individuals rather than grouped into mass classes.

To determine if the slope and intercept of the 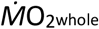-DBA relationship was mass dependent, a LME model was applied (lme4 package; Bates et al., 2015). In this model, VeDBA, mass class, and an interaction between VeDBA and mass class were included as covariates to explain variation in 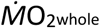 Importantly, inclusion of mass as either a continuous or categorical variable was considered. By including mass class as a continuous variable, the model estimated a single slope or intercept value across all classes, whereas when mass was included as a categorical variable, the model estimated separate slopes or intercepts for each class. Thus, inclusion of mass as a continuous variable in the best-fit model indicated a multiplicative effect of mass on the slope or intercept term, whereas inclusion of mass as a categorical variable indicated an allometric effect of mass on the slope or intercept term. The MuMIn package (Barton and Barton, 2019) was used to build models using every combination of explanatory variables and model fit was compared based on Akaike information criteria (AIC), with decreases in AIC ≥2 considered as improvements in fit (Zuur et al., 2009). Models were fit with a Gaussian error structure, and the final model was validated based on examination of its residuals.

### Allometric scaling of slope and intercept

Following the formation of the best-fit LME model, the allometric scaling rates of the slope and intercept were extracted from the best-fit LME output for each mass class. The estimates and the body mass for each class were natural log-transformed, and a regression analysis was used to determine if there was a significant relationship of *ln*(intercept) and *ln*(slope) with *ln*(mass) (Fig. 2). Lastly, the *ln*(intercept)-*ln*(mass) and *ln*(slope)-ln(mass) regression equations were exponentiated to establish the power functions that described the mass-scale dependence of 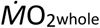-DBA slope and intercept (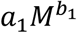 and 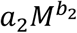 in Equation 6), which we used to build a final predictive model.

**Fig. 2.**
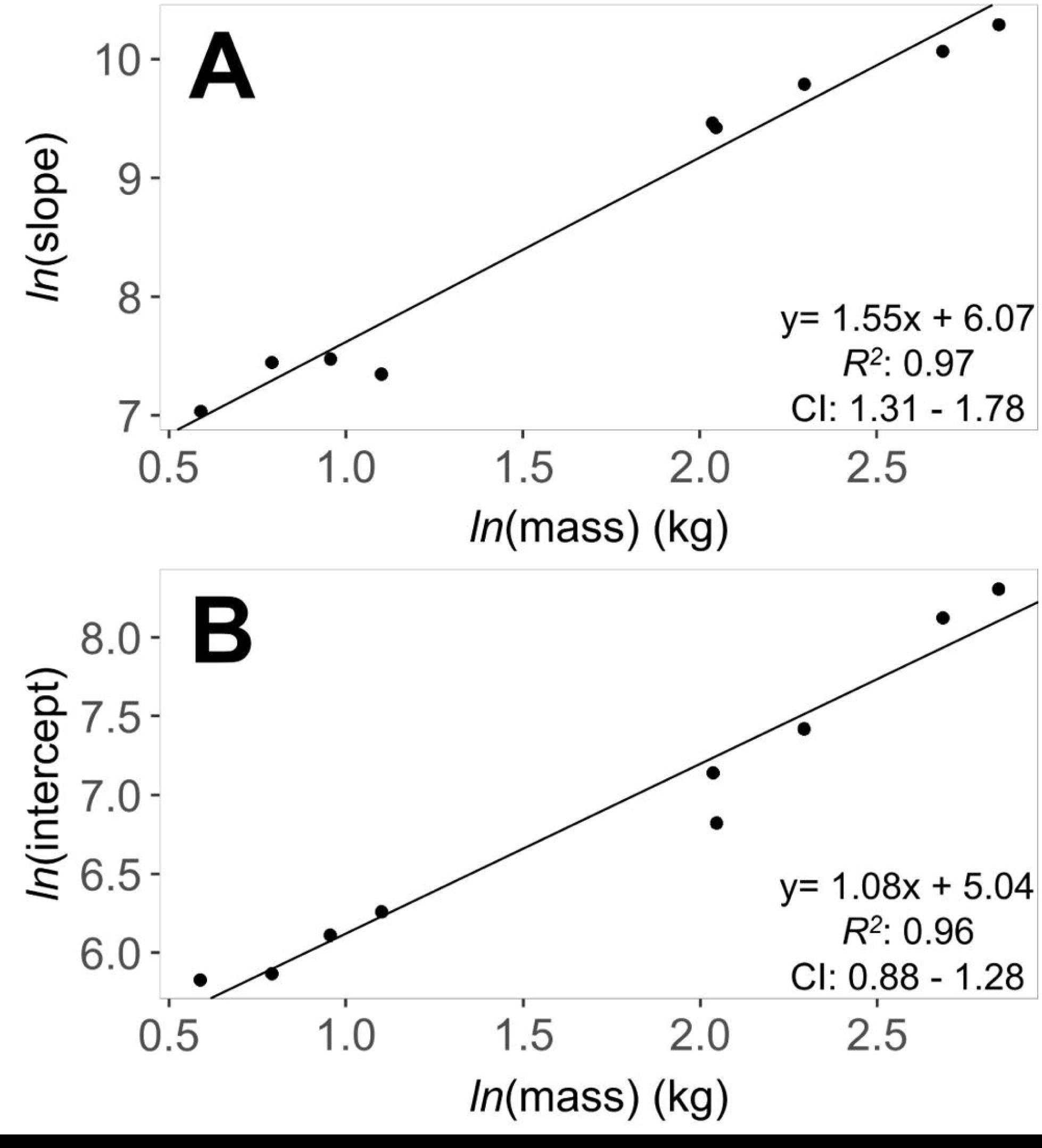
Body mass-scaling of the slope (A) and intercept (B) of the 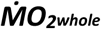-VeDBA relationship. Slope and intercept values were extracted from the best-fit linear mixed-effects model, natural-log transformed, and regressed against the natural-log of body mass. Linear regression equations and associated correlation coefficient (*R*^*2*^) and slope confidence intervals (CI) are displayed for each scaling relationship.

### Model validation

A bootstrap approach was used to validate our final 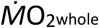 predictive model. 1000 pseudo-datasets were produced via bootstrapping, and for each bootstrapped dataset, a new predictive model was established by applying the same modelling process as above (sections 2.3.2 & 2.3.3). The predictive models were then used to estimate 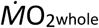 for each observation of VeDBA within the respective dataset. The root-mean square (RMS): 

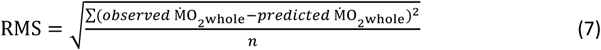

and coefficient of variance (COV): 

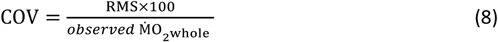

were calculated for each bootstrapped model. To assess the error associated with inactive versus active periods, RMS and COV were also calculated based on 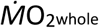 estimations from only inactive and active data, respectively. The overall, inactive, and active mean RMS and COV were calculated to represent model error. In addition, the *R*^*2*^ was extracted from each model and the overall mean used as another quantification of 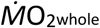 predictive model performance.

### Model comparison: Coefficient-corrected vs. response corrected

Commonly, mass-specific 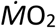(mg O_2_ kg^-1^ hr^-1^) is used when modelling the 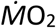-DBA relationship, rather than modelling whole animal 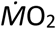(mg O_2_ hr^-1^). By using mass-specific 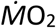, these methods incorporate mass-corrections into the response variable of predictive equations, as opposed to incorporating mass-corrections into the covariates of predictive models, as was done here. To compare the modelling approach developed here with these ‘response-corrected’ approaches (Fig. 1), two additional models were fit to mass-specific 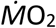 data. For the first model, 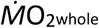 estimates were isometrically corrected by dividing 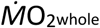 by animal mass in kg (*M*), resulting in mass-specific 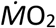 in mg O_2_ kg^-1^ hr^-1^. Secondly, 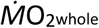 estimates were allometrically corrected by dividing 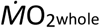 by animal mass in kg raised to an allometric SMR/BMR scaling exponent (*M*^*b*^), resulting in mass-specific 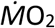 in mg O_2_ kg^-b^ hr^-1^. For this study, an allometric exponent of 0.86 was used, a value widely regarded as the general elasmobranch SMR mass-scaling exponent (Sims, 2000). Models were fit using the glm function, assuming a Gaussian error structure. These response-corrected models will be hereafter referred to as the ‘isometric model’ and ‘allometric model’, whereas the two-step modelling process established in herein will be referred to as the ‘covariate-corrected model’.

To allow comparison between all three modelling approaches, mass-specific estimates from the isometric and allometric models were corrected to 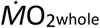 by multiplying estimates by *M* and *M*^*0.86*^, respectively.

To explore estimation error as a function of body mass and activity level, we repeatedly simulated a day in the life of 800 sharks, 100 per mass class. A pseudo-observation was created every 15 minutes over a 24-hour period for each shark by resampling raw observations from sharks within the same mass class via bootstrapping. Before bootstrapping, raw observations were split into two resampling pools, separating inactive and active raw observations. Pseudo-observations were then bootstrapped from these resampling data pools. Inactive pseudo-observations were created by sampling from the inactive data pool, whereas active pseudo-observations were created from the active data pool. This process was repeated for each shark, varying the amount of time sharks spent active between 0–100%, increasing by one percentage-point for each successive iteration. This resulted in 96 pseudo-observations per shark within each iteration. 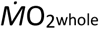 predictive models were established in each iteration using the appropriate predictive equation from the covariate-corrected, isometric, and allometric modelling techniques, and RMS and COV were calculated for each model. 12.5 kg and 14.65 kg mass classes were excluded from the simulation, as both inactive and active data were not available for these classes.

## RESULTS

### Sample size

A total of 43 (mean=1.95 per individual fish, s.d.=0.88) respirometry trials were conducted on 21 individuals, ranging in body mass from 1.74 to 17.15 kg (mean=4.81 kg, S.D.=4.55; Table S1). Once grouped by mass, a total of 10 individual mass classes were retained (Table S1). From these trials, 129 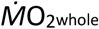 estimations (mean per class=12.90, S.D.=10.23) were obtained for calibrations. However, five 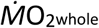 observations were deemed as biologically impossible because they were twice as high as other estimates with similar VeDBA and were removed from analysis, leaving 124 observations (mean per class=12.40, S.D.=10.21), including 37 inactive (mean per class=3.70, S.D.=5.85) and 87 active (mean per class=8.70, S.D.=8.66) (Table S1). From these 124 observations, 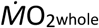 ranged from 286.57 to 6493.68 mg O_2_ hr^-1^ (Fig. 3, Table S1). After correction for acceleration sensor noise (0.018 g), VeDBA ranged from -0.014 to 0.172 g (Fig. 3). Water temperature during calibration intervals ranged from 27.32 to 31.42 °C (mean=29.52 °C, S.D.=0.81) (Table S1).

**Fig. 3.**
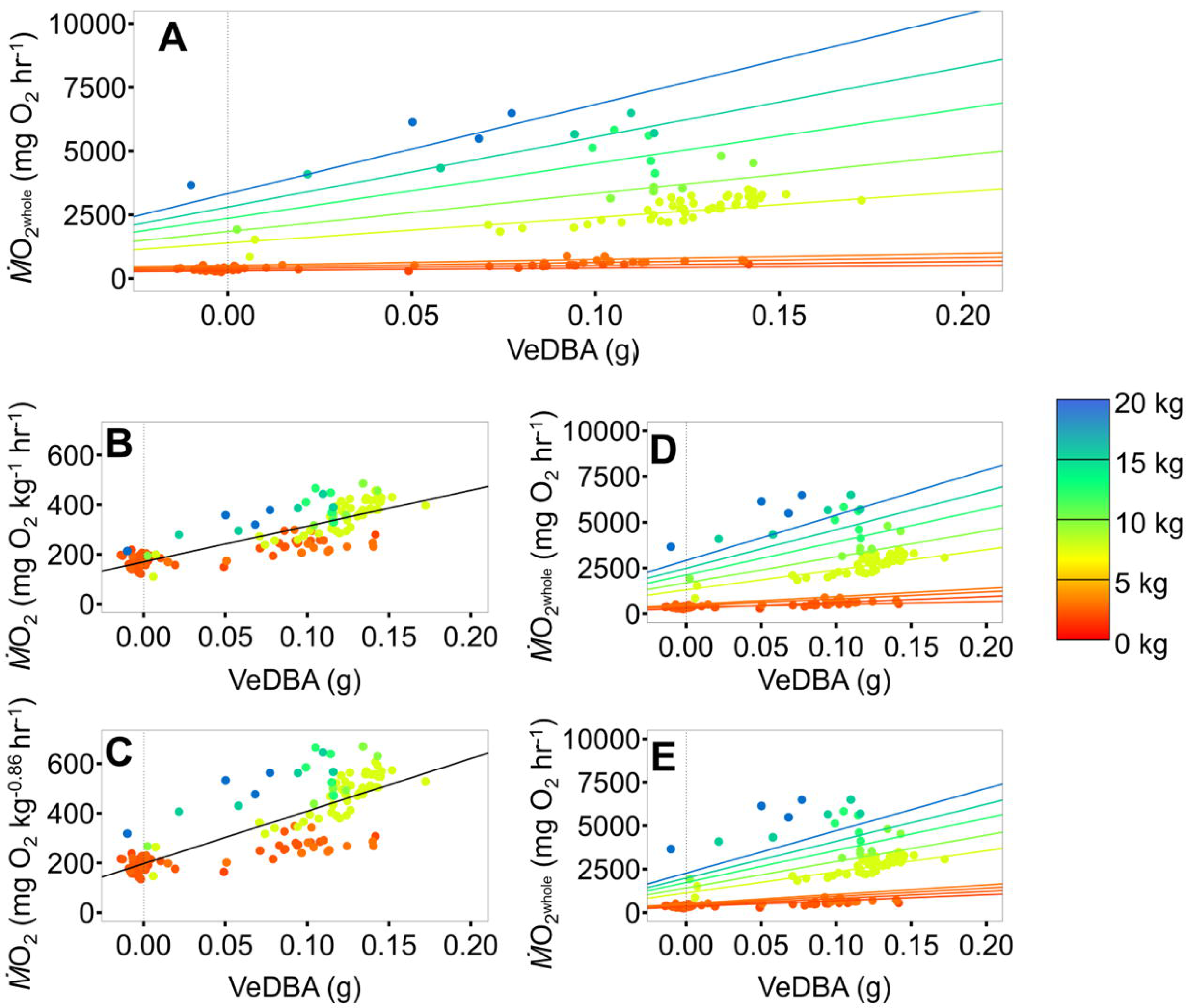
Model predictions overlaid on observed data from respirometry calibration trials. The covariate-corrected model (A) was fit to 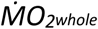 values, whereas the isometric model (B) and allometric model (C) were fit to mass-specific 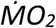 values. For comparison between models, isometric and allometric fits were corrected to 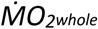 by multiplying estimates by a factor of M^1^ (D) and M^0.86^ (E), respectively. Observations and linear model fits are colour coded according to mass class. Vertical dotted lines represent the point of no activity (y-axis), indicating the intercept of linear estimators

### Final covariate-corrected model and validation

There was a strong positive correlation between VeDBA and 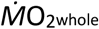, with the slope and intercept displaying significant allometry (Fig. 3). The best-fit model included VeDBA and an effect of mass class as a categorical variable on both the slope and intercept terms (Table 1). From this model, a total of nine slopes and intercepts were extracted from the linear fits for each mass class to assess the mass-scaling of the 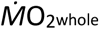-VeDBA relationship. The 12.5 kg mass class was excluded from this scaling assessment because all 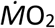 observations for this class occurred over a small range of DBA values, which resulted in a negative linear relationship (Fig. S1). The intercept of the 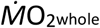-VeDBA relationship increased with a mass-scaling exponent of 1.08 (CI=0.88–1.28, *R*^*2*^=0.96, Fig. 2A), whereas the slope increased with a mass-scaling exponent of 1.55 (CI=1.31–1.78, *R*^*2*^=0.97; Fig. 2B), yielding a final predictive equation of:

**Table 1.** Model selection table for the covariate-corrected linear mixed-effects model. This model was for the first step of the modelling approach (section 2.3.2) developed in this study. Mass was included as both a continuous variable (Mass_cont_) or as a categorical variable (Mass_cat_). * indicates an interaction between terms. Models are ranked based on Akaike information criteria (AIC) values. Degrees of freedom (df) and log-likelihood (logLik) are shown for each model.

| Model | Formula | df | logLik | AIC | ΔAIC | Model weight |
| --- | --- | --- | --- | --- | --- | --- |
| 1 | VeDBA*Mass<sub>cat</sub> + Mass<sub>cat</sub> | 21 | -847.2 | 1736.5 | 0 | 0.964 |
| 2 | VeDBA*Mass<sub>cont</sub> + Mass<sub>cat</sub> | 13 | -857.4 | 1740.8 | 4.38 | 0.036 |
| 3 | VeDBA*Mass<sub>cont</sub> + Mass<sub>cont</sub> | 5 | -888.5 | 1786.9 | 50.45 | 0 |
| 4 | VeDBA + Mass<sub>cat</sub> | 12 | -908.3 | 1840.5 | 104.07 | 0 |
| 5 | VeDBA + Mass<sub>cont</sub> | 4 | -934 | 1876.1 | 139.64 | 0 |
| 6 | Mass<sub>cat</sub> | 11 | -932.3 | 1886.6 | 150.1 | 0 |
| 7 | Mass<sub>cont</sub> | 3 | -955.7 | 1917.4 | 180.98 | 0 |
| 8 | VeDBA | 3 | -1075 | 2156.9 | 420.43 | 0 |
| 9 | Intercept | 2 | -1097 | 2198.5 | 462.02 | 0 |

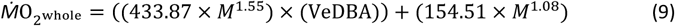

where *M* is individual wet body mass in kg.

The final predictive model explained 98.23% of the variance (*R*^*2*^) in 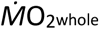 Overall, the model tended to overestimate 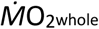 whole, with a mean error (COV) of 19.54% (RMS=329.99 mg O_2_ hr^-1^). When only considering active data, the model had lower error of (COV=17.07%, RMS=376.41 mg O_2_ hr^-1^) than when only considering inactive data (COV=31.65%, RMS=78.34 mg O_2_ hr^-1^).

### Comparison with response-corrected modelling approaches Response-corrected model fits

VeDBA was a significant predictor for both isometrically corrected mass-specific 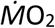(*t*=16.01, *P*<0.001, *R*^*2*^=0.68) and allometrically corrected mass-specific 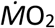(*t*=13.81, *P*<0.001, *R*^*2*^=0.61). The resulting predictive equations were 

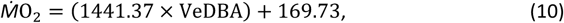

for the isometric model and 

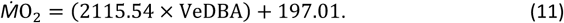

for the allometric model. The isometric model had a mean error of 24.23% (RMS=589.78 mg O_2_ hr^-1^), whereas the allometric model had a mean error of 31.77% (RMS=589.78 mg O_2_ hr^-1^).

### Model error comparison

All models predicted similar 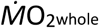 for individuals near the midrange of masses, however, there was greater variation in model estimates for the larger and smaller masses (Fig. 4). For sharks smaller than 7.65 kg, the covariate-corrected model consistently produced the lowest 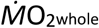 estimates across all activity levels, whereas the allometric model consistently produced the highest 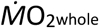 estimates (Fig. 4, S2). On the other hand, for sharks larger than 7.65 kg, the coefficient-corrected model produced the highest 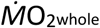 estimates across all activity levels and the allometric model produced the lowest 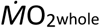 estimates (Fig. 4, S2).

**Fig 4.**
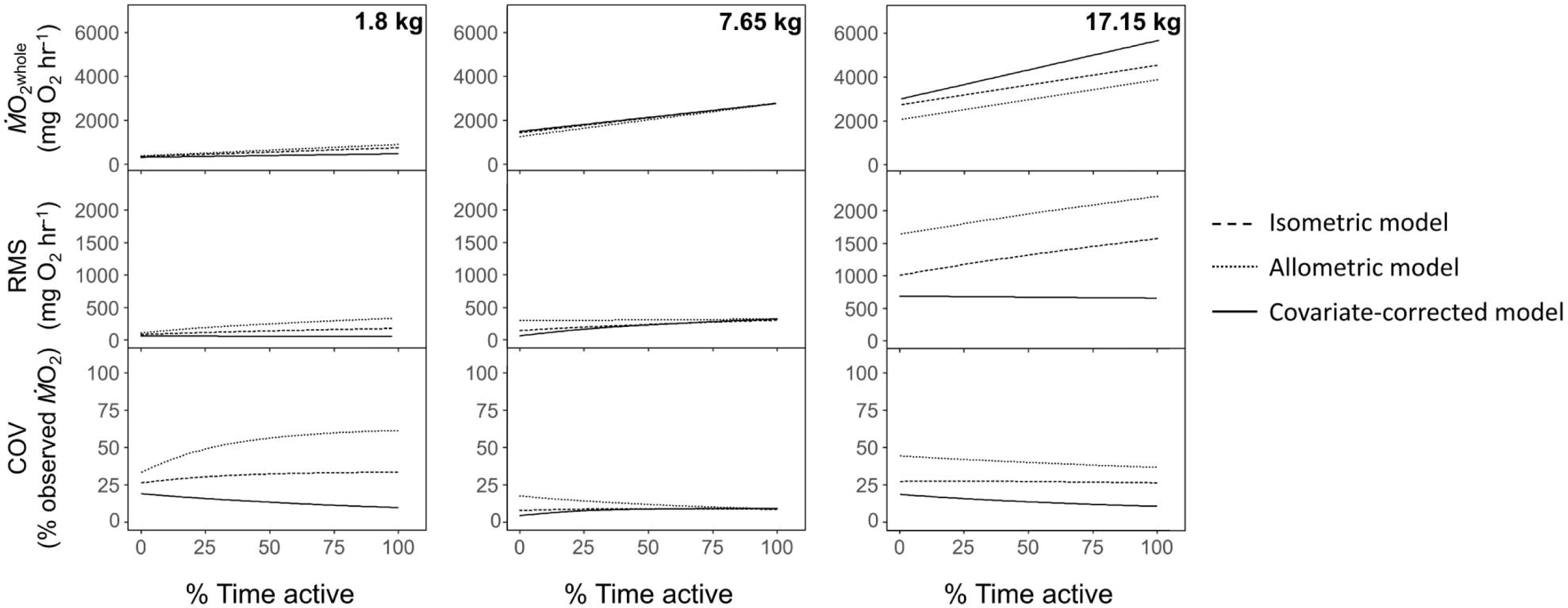
Simulation showing the effect of different modelling techniques when accounting for body mass in 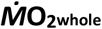 estimations across a range of activity levels. A day in the life of 100 sharks for each mass class was repeatedly simulated, where the proportion of time spent active was varied. For clarity, only the 1.8 kg (left), 7.65 kg (middle column), and 17.15 kg (right) mass classes are shown. Whole animal oxygen consumption rate (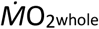; top), root-mean squared error (RMS; middle row), and coefficient of variation (COV; bottom) were calculated over the full range of activity for each body mass using the isometric (dashed line), allometric (dotted line), and covariate-corrected (solid line) modelling approaches.

Overall, the covariate-corrected model consistently had the lowest error across all body sizes and activity levels, whereas the allometric model consistently had the highest error (Fig. 4). The varied activity level simulation revealed systematic biases in the isometric and allometric models that caused estimation error to increase as a function of body mass, whereas no mass-associated bias was present in the covariate-corrected estimates (Fig. 4, S2). Differences in COV between models were smallest for the middle mass classes but varied more widely between models as a function of activity at the lower and higher masses (Fig. 4). At lower masses, the difference between model COV increased with increased time spent active, whereas the difference between model COV at larger masses was relatively consistent (Fig. 4). Overall model COV tended to decrease with increased time spent active, particularly in the covariate-corrected model (Fig. 4).

## DISCUSSION

Despite the prominent influence body size has on many aspects of animal biomechanics and physiology, no studies have tried to describe how the relationship between 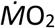 and DBA changes as a function of body size. By individually assessing the scaling of the 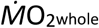-VeDBA slope and intercept, we validated that DBA per unit metabolism is scale dependent, and this dependency varies from that of SMR/BMR. As such, the covariate-corrected modelling approach developed here, which accounts for different body-mass scaling effects within model coefficients consistently had substantially lower error than commonly applied response-corrected models. Ultimately, at the smallest body size, the covariate-corrected model had up to 70.1% and 83.7% lower estimation error than the isometric and allometric model, respectively. When extending to animals in the field, the difference in estimation error between models has large implications for bioenergetics applications. For example, for a 17.15kg lemon shark that is active 50%, the difference between the daily energy expenditure estimated by the isometric model and covariate corrected model would be 17,114.83 mg O_2_ day^-1^. This difference converts to 232.591 kJ day^-1^ (Jobling, 1995). Given that yellow fin mojarra (*Gerres cinereus*), the primary prey of juvenile lemon sharks in Bimini have an average mass of 41.38 g and energy density of 15.58 kJ g^-1^ (Pettitt-Wade et al., 2011), this 232.591 kJ difference represents 0.96 mojarras, or a 25% increase in the expected daily energetic requirement of a shark. With such clear improvements over other modelling techniques, we strongly recommend the covariate-corrected modelling approach for establishing 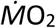 predictive equations across different body sizes.

Although the covariate-corrected approach provides more robust estimates of energy expenditure than currently used techniques, several opportunities for improvement remain. Foremost, while conducted pre-hoc in this study, an ideal full model should incorporate a temperature correction factor in the intercept term. Additionally, more balanced sampling across activity levels and mass classes would allow for a mechanistic examination of error accumulation, which may help to identify and incorporate appropriate correction factors into models. Of course, conducting respirometry experiments over a range of body sizes presents logistical complexities that may limit sample size, as was the case in this study. However, in respirometers with small system to animal volume ratios, which facilitate rapid respiration measurement response times (Clark et al., 2013), calibration interval durations could be decreased to facilitate larger sample sizes per trial. While more calibration intervals within trials may also increase variability of sampled activity levels, using forced activity protocols (e.g. swim tunnels or treadmills) would ensure that observations are obtained across all activity levels, although it is of note that such protocols may introduce other bias through altering natural gaits in many species (Lear et al., 2019; Whitney et al., 2018).

Importantly, as the mechanics and external forces defining the 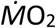-DBA relationship depend on numerous factors such as body form (Alexander, 2005; Bale et al., 2014), locomotory type (Alexander, 2005; Schmidt-Nielsen, 1972), and viscosity of environmental medium (Schmidt-Nielsen, 1972; Wieser and Kaufmann, 1998), 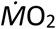-DBA slope and intercept scaling exponents will inevitably vary between taxa. Experiments conducted across taxa will determine the scatter of the scaling exponents and may help identify if laws of biomechanics and physics define universal ratios of slope to intercept scaling exponents, allowing predictive equations to be extrapolated across species.

In circumstances where respiration measurements cannot be acquired from a range of body masses (e.g. facility or animal availability limitations), and thus mass-specific models must be extrapolated, the biases of such models must be carefully considered. Foremost, bias drastically increases with increasing difference in body mass between the median mass of individuals in the calibration experiments and individuals for which respiration rate is being estimated. Thus, to minimize estimation error, caution must be exercised when extrapolating isometrically or allometrically corrected estimates to animals with substantially different body masses than those used in calibration experiments. Additionally, model bias tended to increase with relative activity level, particularly for smaller body masses. This bias may simply be a product of the relatively larger spread of 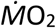 and VeDBA measurements during active calibration intervals compared to inactive intervals. Using forced activity protocols to ensure more balanced sampling of different activity levels throughout calibration experiments would help to elucidate the source of this error. Nevertheless, this activity level associated error indicates that as animals become more active, post-hoc corrections based on isometric or allometric SMR/BMR scaling rates introduce increased bias. Thus, it is imperative to independently account for separate scaling rates of the 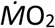-DBA slope and intercept when estimating respiration of highly active species.

We found isometric corrections produced more accurate estimates than using a universal allometric correction of M^0.86^. However, the higher performance of the isometric correction was likely a product of the SMR/BMR scaling rate of lemon sharks in this study being closer to isometry. In species with lower SMR/BMR scaling exponents, like mammals (White and Seymour, 2003), it is likely that a lower nonproportional allometric mass-correction would perform better. However, the higher performance of the isometric correction may also be due to the substantially greater scaling rate of the 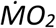-DBA slope than the intercept (i.e. SMR/BMR). Nevertheless, the covariate-corrected modelling approach established herein circumnavigates such issues by separately identifying slope and intercept scaling rates and outperforms commonly applied response-corrected modelling approaches. As such, where possible, the covariate-correction approach should be used for establishing 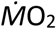-DBA predictive models.

## Acknowledgements

Research was conducted under permits from the Bahamas Department of Marine Resources (MA&MR/FIS/178), Murdoch University Animal Ethics (RW3119/19), and Mote Marine Laboratory Institutional Animal Care and Use Committee (09-09-NW1). We thank Bimini Biological Field Station staff, including C. White, V. Heim, C. Mason, S. Hart, E. Richardson, H. Lintott, D. Warburton, W. Nambu, A. Warrior, K. Yang, and J. Whicheloe, as well as L. Harlow and numerous other interns who were essential for capturing sharks and conducting field trials in Bimini. Mote Marine Laboratory staff, including J. Morris, A. Ontkos, A. Osowski, A. Andres, and R. Hueter, and numerous interns were essential to captive maintenance of sharks and respirometry experiments at MML. Additionally, we thank R. Daly, S.H. Gruber, and T.L. Guttridge for providing logistical support.

## Competing interests

The authors have no conflicts of interest to declare.

## Author contributions

E.E.B. and A.C.G. conceived the study. K.O.L. conducted respirometry trials at Mote Marine Laboratory with help from L.R.B. and N.M.W. E.E.B. conducted trials at Bimini Biological Field Station with help from M.J.S. K.O.L. processed data from Mote Marine Lab trials with A.C.G. and N.M.W. E.E.B. conducted data analysis with help from N.J.A. and A.C.G. All authors contributed to writing the manuscript.

## Funding

Research was generously supported by grants from Save Our Seas Foundation (project no. 402) to E.E.B. and A.C.G and the National Science Foundation (OCE no. 1156141) to N.M.W., A.C.G., and R.E. Hueter. E.E.B. was supported by an Australian Government Research Training Program Scholarship throughout this study.

## Data availability

Data will be deposited in the Dryad Digital Repository if accepted for publication.

